# YSL3-mediated copper distribution is required for fertility, grain yield, and size in *Brachypodium*

**DOI:** 10.1101/2019.12.12.874396

**Authors:** Huajin Sheng, Yulin Jiang, Maryam Rahmati Ishka, Ju-Chen Chia, Tatyana Dokuchayeva, Yana Kavulych, Tetiana-Olena Zavodna, Patrick N. Mendoza, Rong Huang, Louisa M. Smieshka, Arthur R. Woll, Olga I. Terek, Nataliya D. Romanyuk, Yonghong Zhou, Olena K. Vatamaniuk

**Affiliations:** Soil and Crop Sciences Section, School of Integrative Plant Science, Cornell University, Ithaca, NY 14853; Plant Biology Section, School of Integrative Plant Science, Cornell University, Ithaca, NY 14853; Triticeae Research Institute, Sichuan Agricultural University, Wenjiang, Sichuan, China; Cornell Nutrient Analysis Laboratory, School of Integrative Plant Science, Cornell University, Ithaca, NY 14853; Ivan Franko National University of Lviv, Lviv, 79005, Ukraine; Cornell High Energy Synchrotron Source (CHESS), Ithaca, NY 14853

## Abstract

Addressing the looming global food security crisis requires the development of high yielding crops. In this regard, the deficiency for the micronutrient copper in agricultural soils decreases grain yield and significantly impacts a globally important crop, wheat. In cereals, grain yield is determined by inflorescence architecture, flower fertility, grain size and weight. Whether copper is involved in these processes and how it is delivered to the reproductive organs is not well understood. We show that copper deficiency alters not only the grain set but also flower development in both wheat and it’s recognized model, *Brachypodium distachyon*, We then show that a brachypodium yellow-stripe-like 3 (YSL3) transporter localizes to the phloem and mediates copper delivery to flag leaves, anthers and pistils. Failure to deliver copper to these structures in the *ysl3* CRISPR/Cas9 mutant results in delayed flowering, altered inflorescence architecture, reduced floret fertility, grain number, size, and weight. These defects are rescued by copper supplementation and are complemented by the *YSL3* cDNA. This new knowledge will help to devise sustainable approaches for improving grain yield in regions where soil quality is a major obstacle for crop production.

## Introduction

Global food security and the demand for high-yielding grain crops are among the most urgent drivers of modern plant sciences due to the current trend of population growth, extreme weather conditions and decreasing arable land resources [1]. The grain yield is directly linked to the crop and soil fertility. In this regard, it has been known for decades that the deficiency for the micronutrient copper in alkaline, coarse-textured or organic soils that occupy more than 30% of the world arable land, compromises crop fertility, reduces grain/seed yield and in acute cases results in crop failure [2–5]. In accord with the essential role of copper in reproduction, recent studies using synchrotron x-ray fluorescent (SXRF) microscopy established that copper localizes to anthers and pistils of flowers in a model dicotyledonous species, *Arabidopsis thaliana*, and failure to deliver copper to these reproductive organs severely compromises fertility and seed set [6]. Although copper deficiency can be remedied by the application of copper-based fertilizers, this approach is not environmentally friendly and can lead to the build-up of toxic copper levels in soils [2, 5, 7]. Mineral nutrient transporters have been recognized as key targets for improving the mineral use efficiency in sustainable crop production [8]. Wheat is the world’s third important staple crop after maize (*Zea mays*) and rice (*Oryza sativa*); however, wheat grain yield remained relatively low under marginal growing environments despite significant breeding efforts [9]. Wheat is also regarded as the most sensitive to copper deficiency [2, 3, 5]. How copper uptake and internal transport is achieved in wheat and how it affects fertility, is poorly understood. Based on studies in *A. thaliana*, copper uptake and internal distribution is mediated by CTR/COPT transporters, P-type ATPases and members from the Yellow Stripe-Like (YSL) subfamily of the oligopeptide (OPT) transporter family [7, 10–17]. The majority of these transporters are transcriptionally upregulated by copper deficiency by a conserved transcription factor, SPL7 (**S**quamosa **P**romoter Binding Protein–**l**ike7), and a newly identified transcription factor CITF1 (**C**opper-Deficiency **I**nduced **T**ranscription **F**actor1) [6, 18, 19].The expression of several COPT family members is also induced in roots by the copper deficiency in *Oryza sativa* and an emerging wheat model *Brachypodium distachyon* (from here on brachypodium), and several brachypodium COPTs mediate low-affinity copper uptake [20, 21]. A member of the YSL transporters, OsYSL16 functions in the phloem-based copper delivery to reproductive organs in rice [22–24]. Other studies, however, reported that OsYSL16 functions mainly in the distribution of iron [25, 26]. Recognizing the limitations of wheat for functional genetics studies due to polyploidy, lower transformation rates and longer life cycle, we used brachypodium as a wheat proxy [27–30] for the study of copper transport processes and their role in establishing yield traits. We show that copper deficiency alters not only the grain set but also flower development in both wheat and brachypodium. We reveal that brachypodium yellow-stripe-like 3 (YSL3) transporter mediates phloem-based copper distribution from mature leaves to flag leaves, anthers and pistils of florets. Loss of this function in the *ysl3* mutant results in a delayed flowering, altered inflorescence architecture, reduced floret fertility, grain number, size, and weight. These defects are rescued by copper supplementation and are complemented by the *YSL3* cDNA. Our results suggest that the manipulation of YSL3 and other-like proteins has the potential to play a role in devising sustainable and environmentally friendly approaches for improving wheat and other cereal grain yields and thus, food security.

## RESULTS

### Copper Deficiency Significantly Decreases Flower Formation and Seed Yield in Wheat and Brachypodium

We first evaluated the growth and fertility of wheat and brachypodium grown under different concentrations of copper to validate using brachypodium as a wheat mode in this study. Omitting copper from the hydroponic medium severely stunted the growth, tiller, head, flower and seed/grain formation per plant in both wheat and brachypodium (**Fig. 1** and **Supplemental Fig. 1**). Low copper (10 nM) while reduced the number of tillers and heads per plant of both plant species (**Supplemental Fig. 1C, D, E, F),** exerted the most pronounced effect on flower and seed formation (**Fig. 1E** to **H**). Notably, seed formation was reduced by 87% in both wheat and brachypodium when plants were grown under 50 nM copper, although flower formation was only somewhat reduced compared to plants grown under copper replete conditions (125, 250 nM copper, **Fig. 1E** to **H**). These data show that copper deficiency impacts different aspects of reproductive development including flower and seed/grain formation, with the most dramatic effect on seeds/grain production. Furthermore, these data supported the applicability of using brachypodium for the study of the relationship between copper and fertility in cereals as well as the identification of transport pathways responsible for the delivery of copper to plant reproductive organs.

**Figure 1.**
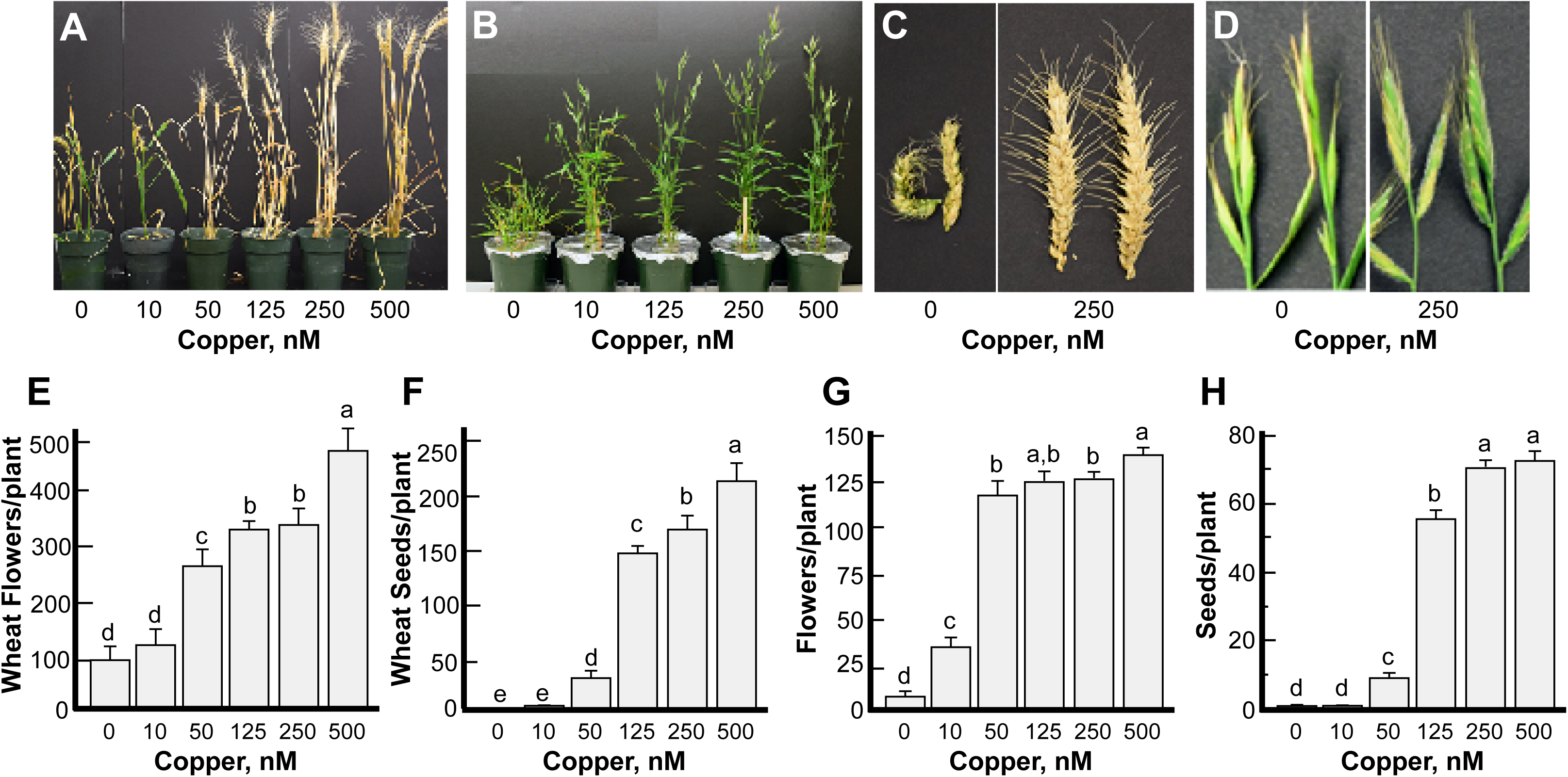
Copper deficiency alters flower development and reduces grain number in wheat and brachypodium. Plants were grown hydroponically under indicated copper concentrations. (**A** to **D)** show representative images of plants, tiller and head appearance under different copper concentrations of wheat (**A, C**) and brachypodium (**B, D**). (**E)** to (**H)** show the number of flowers and grains per plant in wheat (**E, F**) or brachypodium (**G, H**). Statistical analysis in (**E)** to (**H)** was done with one-way ANOVA in the JMP Pro 14 software package; comparison of the means for each pair was done using Student’s *t*-test. **E** to **H** show mean values ± S.E. from the analysis of 4 (wheat) to 6 (brachypodium) independently grown plants from one out of two (wheat) and three (brachypodium) independent experiments. Levels not connected by the same letter are significantly different (*p* < 0.05).

### Copper Deficiency Increases the Transcript Abundance of *YSL3*

We then focused on brachypodium YSL3 because its counterparts in *Arabidopsis* and rice contribute to transition metals, including copper transport [15, 22, 23, 28]. We found that *YSL3* was expressed in different plant organs including roots, leaves, nodes and reproductive organs **(Fig. 2A**). The highest expression of Y*SL3* was observed in young leaves of four-week-old seedlings, followed by flag leaves at the flowering stage and mature leaves at jointing (**Fig. 2A**). *YSL3* was also expressed in different flower organs including lemma, palea and ovaries, but the abundance of the transcript was much lower than in leaves (**Fig. 2A**).

**Figure 2.**
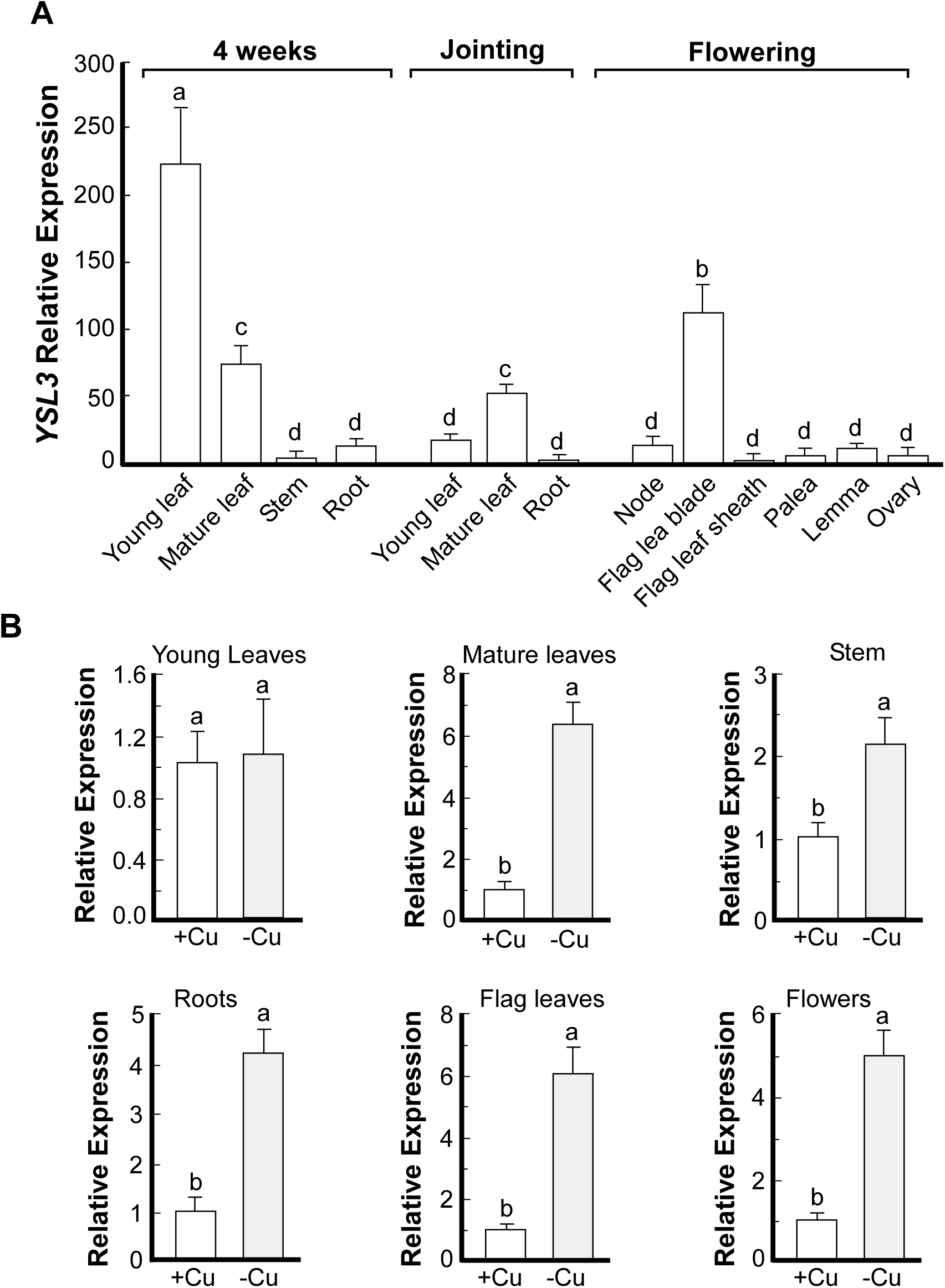
Copper deficiency increases transcript abundance of *YSL3*. **(A)** The expression level of *YSL3* in different tissues at different growth stages. Wild-type *Brachypodium* was grown hydroponically under copper sufficient (0.25 µM CuSO_4_) conditions. The indicated plant tissues were collected at the indicated time and developmental stage; RNA was extracted and subjected to RT-qPCR analysis. (**B**) The expression level of *YSL3* in different tissues of *Brachypodium* wild-type grown hydroponically under copper sufficient (**+Cu**) or deficient conditions (**-Cu**) conditions. Young (2 uppermost leaves) and mature leaves (the remaining leaves), stems and roots were collected from four-week-old plants. Flag leaves and flowers were collected from six-week-old plants. Copper deficiency was achieved by transferring plants to a fresh medium lacking copper one week prior to tissue sampling. **A** and **B** show mean values from 3 independent experiments; error bars show S.E. Tissues were pooled from 3 plants per each experiment. Levels not connected by same letter are significantly different (*p* < 0.05).

We then found that *YSL3* was highly upregulated under copper deficiency in roots, stems and mature leaves but not in young leaves of four-week-old plants. Copper deficiency also significantly increased the transcript abundance of *YSL3* in flag leaves and flowers at the reproductive stage (**Fig. 2B**). These results suggested that YSL3 might be involved in internal copper distribution and delivery to reproductive organs.

### *YSL3* is Expressed Mainly in the Phloem and Localizes to the Plasma Membrane

We next examined the tissue and cell-type specificity of *YSL3* expression using *Brachypodium* transformed with the *YSL3_pro_-GUS* construct. We found that YSL3 is expressed predominantly in the vascular tissues of roots and leaves of plants subjected to copper deficiency (**Fig. 3A, B**). The bulk of GUS staining was associated with the phloem of large and small longitudinal veins as well as in mesophyll parenchyma cells (**Fig. 3C**). Because nodes of grasses are regarded as hubs directing metal distribution [31], we also evaluated *YSL3pro-GUS* activity in the node I. GUS activity was observed mainly in large vascular bundles of the node (**Fig. 3D**). Within the large vascular bundles, xylem is located on the inside and the phloem is on the outside and GUS staining was mainly associated with the phloem and also was found in parenchyma cells (**Fig. 3E**). Concerning florets, GUS activity was observed in the ovary, styles (**Fig. 3F**), the vasculature of the lemma (**Fig. 3G**), but not in anthers and palea. GUS activity was undetectable in any of the tissues of plants grown under copper sufficient conditions. The predominant expression of *YSL3* in the phloem, phloem parenchyma cells and mesophyll suggested that it is involved in internal copper distribution rather than copper uptake into the roots. We next found that YSL3 localizes to the plasma membrane (**Supplemental Figure 2**), suggesting that it is involved in movement of substrates into or out of the cell rather than subcellular (*e.g.* vacuolar) sequestration.

**Figure 3.**
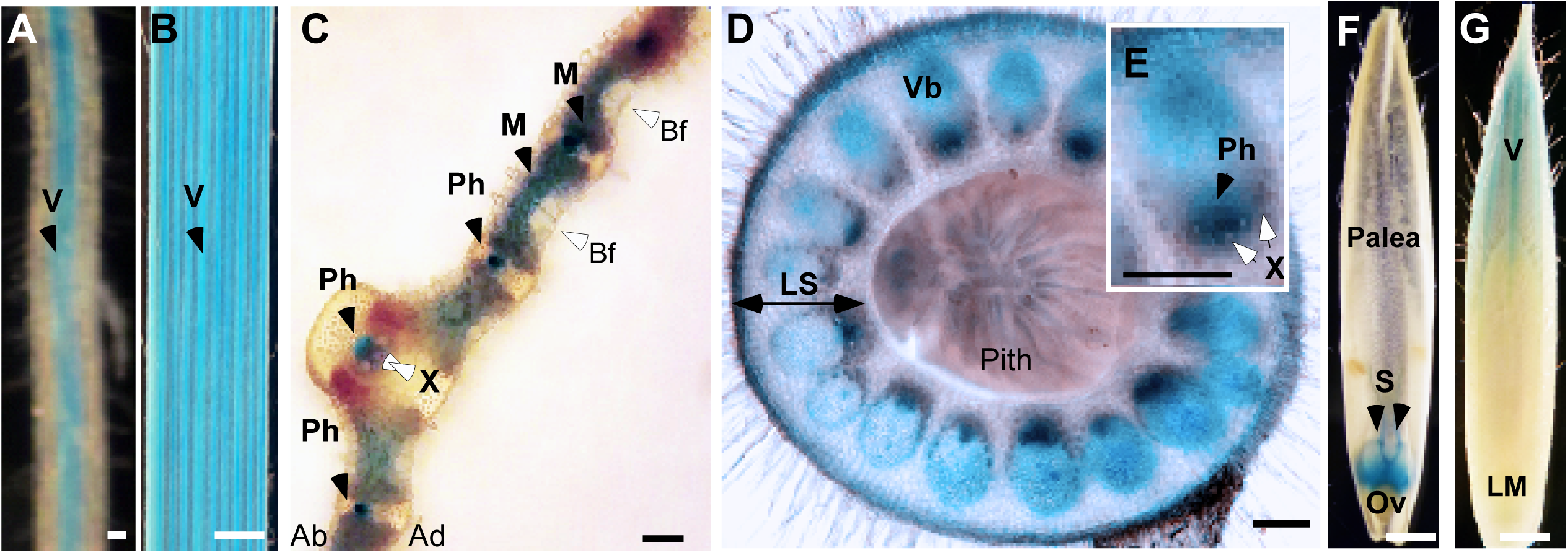
Tissue-specificity of the expression of *YSL3*. Transgenic plants expressing *YSL3_pro_*-GUS construct were grown hydroponically for 3 (**A** to **C**) or 6 (**D** to **G**) weeks. Plants were transferred to hydroponic solution without copper (-Cu) for 1 week prior to histochemical analysis. (**A**) and (**B**) show representative images of GUS staining in the vasculature (**V**, black arrow) of lateral roots and the third emerged leaf, respectively. Transverse sections through a leaf lamina (**C**), and a node (**D**) show GUS staining in tissues indicated by black arrows. (**E**) shows a close-up of a vascular bundle embedded in a dashed box in (**D**). (**F**) shows GUS staining in dissected flowers of *YSL3_pro_*-GUS-expressing transgenics grown under copper deficiency. (**G)** shows GUS staining in the vasculature of the lemma of plants grown under copper deficiency. X-xylem vessels; Ph-phloem, Vb – vascular bundle; LS – leaf sheath of the node, V - vasculature; S – styles of pistils; Ov – ovary; LM – lemma; Bf – bulliform cells, Ad is the adaxial, Ab is the abaxial side of the leaf lamina. *YSL3_pro_-GUS*-mediated staining is indicated by black arrows. Open arrows point to other tissues and cell-types. GUS staining was not detected in plants grown under copper sufficient conditions. Scale bars: 100 µm for **A**, **C**, **D**, **E** and 1 mm for **B**, **F**, **G**.

### The *ysl3-3* Mutant of Brachypodium is Sensitive to Copper Deficiency

We then generated *ysl3* deletion mutants using the CRISPR/CAS9 (clustered regularly interspaced short palindromic repeats) approach (**Supplemental Information** and **Supplemental Figures 3A** and **4)**. After obtaining Cas9-free mutant lines (**Supplemental Figure 3B, C**), positions of deletion breakpoints were established by sequencing. Three alleles bearing 122, 123 and 182 bp deletions encompassing a part of the 5′UTR and the first exon of *YSL3* were identified and designated as *ysl3-1*, *ysl3- 2* and *ysl3-3*, respectively (**Supplemental Information online** and **Supplemental Figures 3A, E)**. Plants of all alleles were smaller than wild-type when were grown under copper sufficient conditions (**Supplemental Figure 3D**). Given the essential role of copper in plant growth and development, we hypothesized that the smaller size of mutants could be due to a defect in the copper transport.

We then used the *ysl3-3* allele, for the in-depth studies and have also generated two *ysl3-3* transgenic lines expressing *YSL3* cDNA, *ysl3/YSL3-1* and *ysl3/YSL3-2,* for functional complementation assays. The level of *YSL3* transcript was increased in both *ysl3/YSL3-1* and *ysl3/YSL3-2* lines compared to the wild-type (**Supplemental Figure 5**). We next compared the growth of the *ysl3-3* mutant *vs*. wild-type and *ysl3/YSL3-1* and *ysl3/YSL3-2* plants in the medium with *vs.* without copper. As evident by the smaller stature of the *ysl3-3* plants (**Supplemental Figure 6A**), curling of their leaf margins (**Supplemental Figure 6B**) and decreased height and dry weight of shoots (**Supplemental Figure 6C, D**), the *ysl3-3* mutant was more sensitive to copper deficiency that the wild-type. The dry weight of roots of the *ysl3* mutant was significantly different from wild-type even when plants were grown under copper sufficiency and omitting copper from the medium did not change it further (**Supplemental Figure 6E**). Importantly, the expression of *YSL3* cDNA in the *ysl3-3* mutant rescued all defects of the mutant (**Supplemental Figure 6A to E**) suggesting that slower growth of the *ysl3-3* plants under control conditions and further reduced growth under copper deficiency were due to the loss of *YSL3* gene. The *ysl3-3* mutant was not more sensitive than wild-type to manganese, iron or zinc deficiencies (**Supplemental Figure 7**). Together, these results indicate that *BdYSL3* is essential for the normal growth of *Brachypodium* under control condition and under copper deficiency.

### The *ysl3* Mutant has a Delayed Flowering Time and Produces more Spikelets and Florets per Inflorescence

We then grew wild-type, the *ysl3-3* mutant and *ysl3/YSL3-1* and *ysl3/YSL3-2* plants in soil to evaluate the role of YSL3 in development and reproduction. We found that the flowering time of the *ysl3-3* mutant was significantly delayed compared to wild-type plants (**Fig. 4A, B**). While wild-type plants have started flowering by the 40^th^ day of growth, the *ysl3-3* mutant flowered on average 2 weeks later (**Fig. 4A, B**). The *ysl3-3* mutant also had shorter flag leaves (**Fig.4C** and **Table 1**) and inflorescences (*alias* spikes) compared to wild-type (**Fig. 4C**). We then compared the flower development of the *ysl3-3* mutant *vs.* wild-type. Florets in grasses are formed on a structure called spikelet. In *Brachypodium*, a terminal spikelet and a limited number of lateral spikelets give rise to a variable number of florets per spikelet [29]. We found that while wild-type plants produced 2 to 4 lateral spikelets in addition to a terminal spikelet, the *ysl3-3* mutant developed 5 to 7 lateral spikelets in addition to a terminal spikelet (**Fig. 4C** and **Table 1**). The increased number of spikelets in the *ysl3-3* mutant resulted in a 1.8-fold increase in the floret number compared to wild-type plants (**Table 1**). Fertilizing the *ysl3-3* mutant with 25 µM CuSO_4_ functionally complemented the mutant suggesting that the decreased flag leaf length, altered spikelet and floret formation in the mutant was due to a defect in copper transport (**Table 1**). Furthermore, the expression of *YSL3* cDNA in the *ysl3-3* mutant also functionally complemented the mutant (**Fig. 4A** to **C** and **Table 1**) suggesting that the decreased flag leaf length, altered spikelet and floret formation in the mutant was due to the loss of *YSL3*.

**Figure 4.**
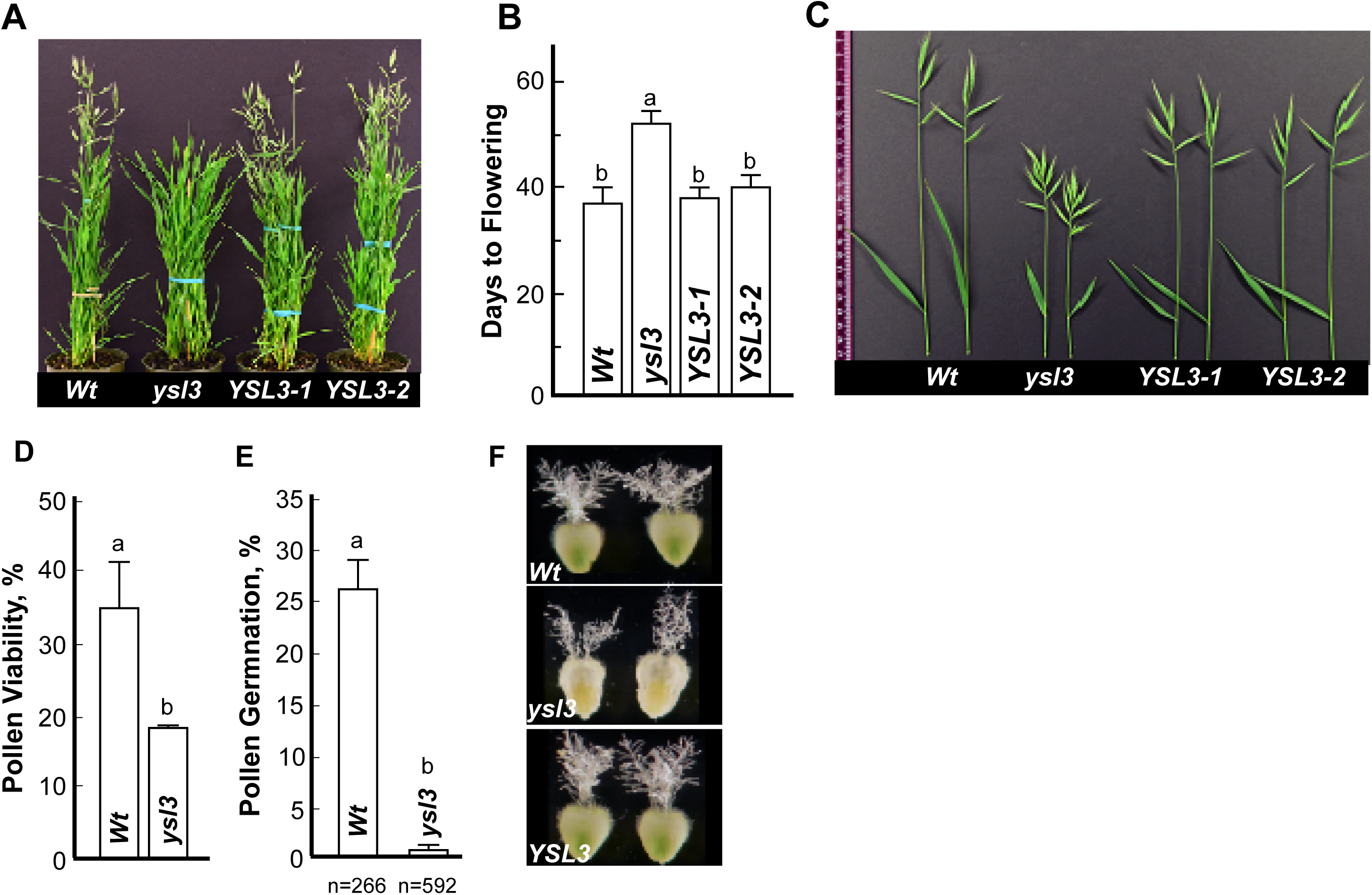
The *ysl3-3* mutant has a delayed flowering time, altered inflorescence architecture and pollen viability. The indicated plant lines were grown in soil and fertilized bi-weekly with the N-P-K fertilizer. (**A**) shows a representative image of the *ysl3-3* mutant which was still in the vegetative stage in contrast to wild-type and *YSL3-1* and *YLS3-2* complementary lines that have reached the reproductive stage of the development. (**B**) shows days to flowering in each of the indicated plant lines. Mean values ± S.E are shown (n = 3 independent experiment with at least 6 plants analyzed per each experiment). Levels not connected by same letter are significantly different (p < 0.05). (**C**) shows a representative image of spikes with a flag leaf collected from plants grown in soil. In order to collect spikes of all plant lines simultaneously, the *ysl3-3* mutant has been germinated two weeks in advance to other plant lines. (**D**) shows the viability of pollen collected from the wild-type (Wt) and the *ysl3-3* mutant (*ysl3*), grown as described above. Mean values of 6 independently grown plants from three independent experiments are shown. Error bars show S.E. Levels not connected by same letter are significantly different (p < 0.05). (**E**) *In vitro* pollen germination assay shows poor germination rate of the *ysl3-3* mutant compared to wild-type. Values are mean ± SE of 4 and 7 independent experiments for wild-type and the *ysl3-3* mutant, respectively; n = number of pollens scored are shown below each bar. Levels not connected by same letter are significantly different (p < 0.01). (**F**) shows the morphology of pistils collected from wild-type, the *ysl3-3* mutant, and the *YSL3-1* complementary line, all grown in soils and fertilized bi-weekly with N-P-K.

**Table 1.** The *ysl3-3* mutant has altered flower morphology and fertility. Plant were grown in soil and fertilized bi-weekly with a standard N-P-K fertilizer. The *ysl3-3* mutant was also grown in soil and in addition to N-P-K was also fertilized bi-weekly with 25 µM CuSO_4_ (*ysl3* +25 µM CuSO_4_). Spikes were collected at the end of reproductive stage. We note that because the *ysl3-3* mutant is developmentally delayed, its spikes were harvested and fertility was analyzed separately although all plant lines were sown and grown concurrently. Mean values ± SE are shown (n = 15 plants per each genotype). Asterisks (*) indicate statistically significant differences from the wild-type (*p* < 0.0001). ^a^Spikelets include the terminal spikelet and lateral spikelets ^b^Florets number indicates the total florets produced per spike ^c^Seeds number indicates the total seeds produced per spike ^d^Fertile florets number was calculated as % of seeds formed per the number of florets per spike

| Genotype | Flag leaf length (cm) | Spikelets<sup>a</sup> | Florets<sup>b</sup> | Seeds<sup>c</sup> | Fertile Florets (%) |
| --- | --- | --- | --- | --- | --- |
| Wt | 7.6 ± 0.23 | 3.8 ± 0.22 | 39.8 ± 3.09 | 29.0 ± 6.71 | 76.2 ± 1.35 |
| *ysl3-3* | 4.9 ± 0.38* | 7.7 ± 0.25* | 70.0 ± 4.56* | 31.0 ± 0.58 | 42.6 ± 3.33* |
| *ysl3/YSL3-1* | 6.4 ± 0.44 | 4.8 ± 0.48 | 41.0 ± 2.52 | 31.3 ± 2.81 | 75.9 ± 3.76 |
| *ysl3* + 25 μM $\text{CuSO}_4$ | 6.7 ± 0.56 | 5.0 ± 8.36 | 46.3 ± 8.36 | 29.0 ± 3.49 | 64.8 ± 4.92 |

### The *ysl3* Mutant has a Defect in Pollen and Floret Fertility

Because the *ysl3-3* mutant developed more florets per plant and spike (**Fig. 4C** and **Table 1**) than wild-type, we anticipated that the mutant would also form more seeds. Surprisingly, there was no difference in grain production per spike between different plant lines (**Table 1**). Furthermore, we found a significant (1.8-fold) reduction in floret fertility as evident by a reduced number of grains formed per the number of florets per spike in the mutant *vs.* wild-type (**Table 1**). Importantly, the expression of *YSL3* in the *ysl3*-*3* mutant or copper supplementation rescued this defect (**Table 1**). We concluded that YSL3-mediated copper delivery to flowers is important for flower fertility. We then examined whether the reduced fertility of the *ysl3-3* mutant is associated with the defect in androecium, gynoecium or both. We found that pollen viability of *ysl3-3* pollen was nearly half-of observed in the wild-type and fewer *ysl3* mutant pollen grains were able to germinate and produce pollen tubes (**Fig. 4D, C**). We also found that more than 40% of the flowers from *ysl3*-*3* mutants had altered stigma morphology compared to the wild-type. Specifically, the stigma of the *ysl3-3* mutant appeared dehydrated, shorter and less feathery compared to the wild-type (**Fig. 4F**). Together, these data suggest that the compromised fertility of the *ysl3*-*3* mutant might be due to defects in both androecium and gynoecium.

### YSL3 Regulates Copper Delivery from Mature Leaves to Flag Leaves and Flowers

To examine whether the delayed transition to flowering and reduced fertility of the *ysl3* mutant were caused by the disruption of copper transport, we analyzed copper concentration and spatial distribution in different plant tissues using inductively coupled plasma mass spectrometry (ICP-MS) and 2D synchrotron-x-ray fluorescence (2D-SXRF) microscopy, respectively. We did not find a significant difference in copper concentration in roots of the *ysl3-3* mutant compared to wild-type or *ysl3/YSL3-1* plants (**Fig. 5A**). However, the *ysl3-3* mutant accumulated 54% more copper in mature leaves compared to wild-type (**Fig. 5B**). The expression of *YSL3* cDNA in the *ysl3-3* mutant reduced copper accumulation in mature leaves to the wild-type level suggesting that the observed defects in the *ysl3*-*3* mutant were due to the loss of YSL3 function (**Fig. 5B**). In contrast to mature leaves, flag leaves and flowers of the *ysl3-3* mutant accumulated 2.9- and 2.6-fold less copper, respectively than corresponding organs of wild-type (**Fig. 5C**). The expression of *YSL3* in the *ysl3* mutant rescued its copper accumulation defect. Together, these results suggested that YSL3 directs copper distribution from mature leaves to flag leaves and flowers.

**Figure 5.**
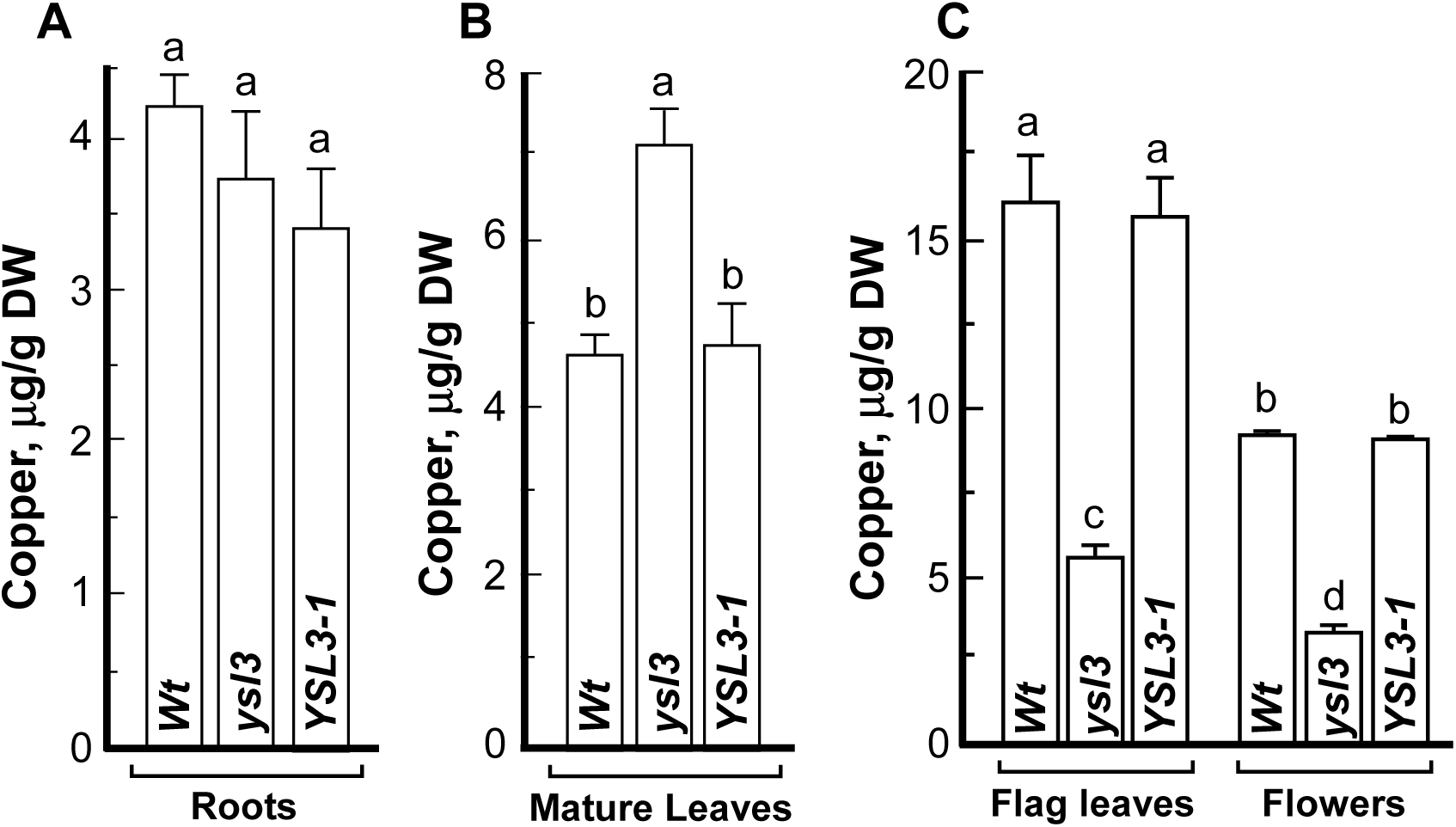
Copper delivery to flag leaves and flowers is impaired in the *ysl3* mutant. ICP-MS- based analysis of the concentration of copper in roots (**A**), mature leaves (**B**), flag leaves and flowers (**C**) of wild-type plants (**Wt**), the *ysl3-3* mutant (***ysl3***) and the *ysl3-3* mutant expressing the *YSL3* cDNA (***YSL3-1***). **A** to **C** show mean values of 3 independent experiments, error bars show S.E. Levels not connected by the same letter are significantly different (*p* < 0.05).

Analysis of mature leaves using 2D-SXRF disclosed that copper was associated mainly with leaf veins in both wild-type and the *ysl3-3* mutant. We also found that copper accumulation was much higher in veins of the *ysl3-3* mutant compared to wild-type (**Fig. 6A**). Similar to mature leaves, the bulk of copper was associated with major and minor veins of flag leaves in wild-type and the *ysl3-3* mutant (**Fig. 6B**). However, both vein types in flag leaves of the *ysl3-3* mutant accumulated much less copper compared to wild-type and the mutant expressing *YSL3* cDNA (**Fig. 6B**). The bulk of copper was associated with anthers and ovary of florets in wild-type while copper was barely detectible in anthers and was significantly lower in the ovary of the *ysl3-3* mutant compared to wild-type and the *ysl3-3* mutant expressing *YSL3* cDNA (**Fig. 6C**).

**Figure 6.**
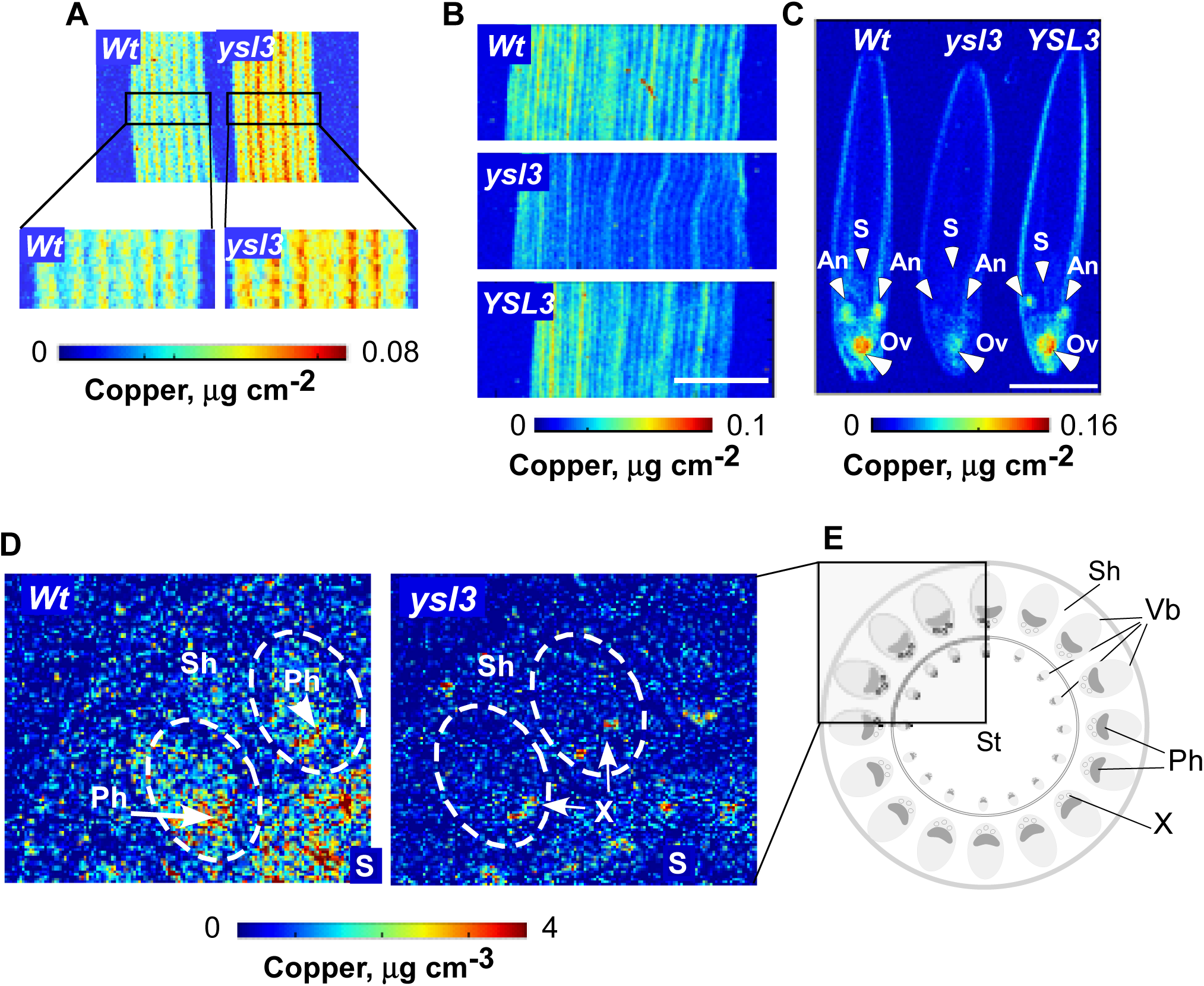
The distribution of copper is altered in the *ysl3-3* mutant. SXRF-based analysis of the spatial distribution of copper in mature leaves (**A**), flag leaves (**B**) and florets (**C**) of the indicated plant lines. Middle part of the leaf was used for SXRF imaging in both **A** and **B**. White arrows in **C** point to anthers (**An**), stigma (**S**) and ovaries (**Ov**). Scale bar = 2 mm. Mature leaves in (**A**) were collected from 3-week-old plants, grown hydroponically with 0.25 µM CuSO_4_. Flag leaves and florets were collected from soil-grown plants that were fertilized bi-weekly with N-P-K. (**D**) Two-dimensional confocal SXRF (2D-CSXRF) was used to visualize the spatial distribution of copper in node I of indicated plant lines. (**E**) illustrates the anatomy of the upper part of the node I. Part of the node in a rectangle was scanned using C-SXRF and is shown in **D**. Sh – leaf sheath, Vb- vascular bundle, Ph – phloem, X- xylem.

We next thought to determine the spatial distribution of copper in nodes because nodes of grasses act as hubs directing and connecting mineral transport pathways for their subsequent distribution to various organs [31]. To do so, we utilized 2D-SXRF in a confocal mode (2D-CXRF) using a specialized x-ray collection optic to obtain micron-scale resolution [32, 33]. For the current study, this technique is preferable to traditional SXRF methods (both 2D SXRF and 3D micro-XRF tomography) because it allows comparison of quantitative metal distributions among different samples without the need to control or limit sample thickness or lateral size [33]. We found that the bulk of copper was associated with large vascular bundles with a higher concentration in the phloem region in nodes of the wild-type (**Fig. 6D, E**). In contrast, copper accumulation in vascular bundles was barely detectible in the *ysl3-3* mutant and was mostly associated with the xylem (**Fig. 6D, E**).

Taken together, these data suggested that *YSL3* plays an important role in copper delivery from mature to flag leaves and then after to reproductive organs and acts by loading copper to the phloem. This YSL3 function is important for the normal development of flowers and fertility.

### Grains of the *ysl3-3* Mutant Accumulate less Copper, are Smaller and Lighter

We next tested whether the loss of the YSL3 function also impacts copper accumulation in grains. We found that the concentration of copper in grains of the *ysl3-3* mutant was lower by 44.52 % compared to wild-type and the *YSL3*-*1* complementary line, all grown in soil (**Fig. 7A**). This shows that the *ysl3-3* mutant defect in remobilizing copper from mature leaves to flag leaves reduces not only copper accumulation in reproductive organs (**Figs. 6B, C**) but also loading to grains.

**Figure 7.**
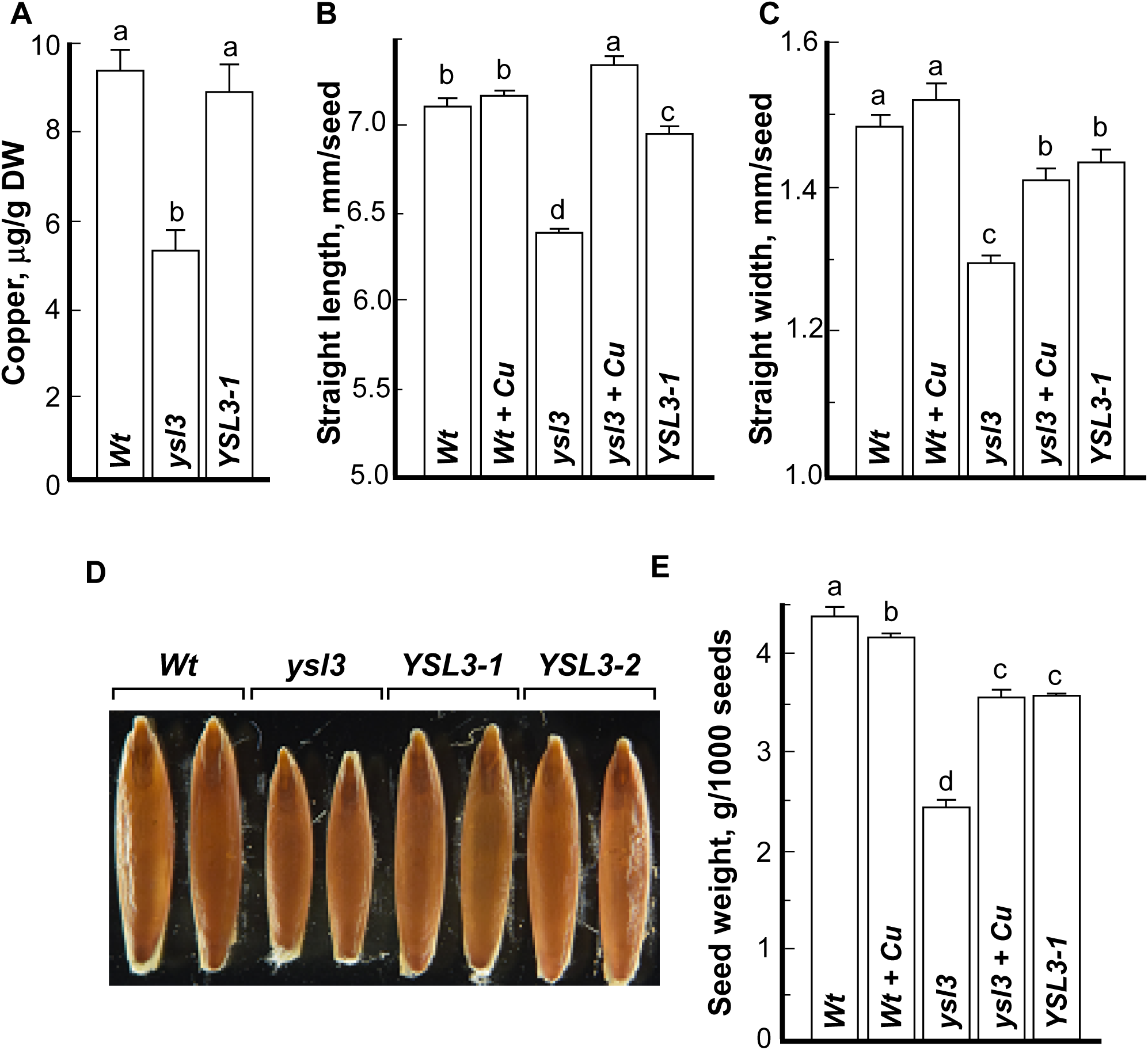
Seeds of the ysl3-3 mutant accumulate less copper, are smaller and lighter. Grains were collected from soil-grown plants that were fertilized bi-weekly with N-P-K. (**A**) ICP-MS analysis of copper concentration in seeds of the indicated plant lines. **B**, **C** shows straight length and width, respectively of grains collected from the indicated plant lines. Grains were dehusked and the straight length and width of randomly selected grains was measured using the WinSEEDLETM of STD4800 Scanner (Regent Instruments Inc., Canada, 2015). (**D**) shows a representative image of seeds pooled from at least three plants from each independent experiment (n=3). (**E**) shows the weight of 1000 dehusked grains of indicated plant lines. Presented values are arithmetic means ± S.E. (n = 3 pools of seeds from five plants per each line; a representative result from 3 independent experimental set-ups is shown). Seeds were pooled together from five plants per line. Levels not connected by same letter are significantly different (p < 0.05).

While dehusking grains of different plant lines for ICP-MS analysis, we noticed that the *ysl3-3* mutant produced shorter and thinner grains than wild-type plants and both complementary lines. This observation was then confirmed by analysis of the straight grain length and width (**Fig. 7B** to **D**). Consistent with a shorter and thinner size, the 1000-grain weight of the *ysl3* mutant was reduced by 30% compared to wild-type (**Fig. 7E**). The expression of *YSL3* cDNA in the *ysl3-3* mutant or copper supplementation rescued the grain size and 1000-grain weight of the *ysl3-3* mutant. These results show that YSL3 function and copper are important for the expression of important agronomic traits including grain size and weight.

## Discussion

Providing a sufficient amount of high quality, nutrient-dense and toxin-free food using sustainable and environmentally friendly approaches are among the grand challenges of the 21^st^ century, driven by the population growth, increasing instances of extreme weather conditions and decreasing arable land resources that limit crop yields [1, 34, 35]. Considering that a micronutrient copper is among yield-limiting factors of a globally important crop, wheat, here we sought to determine how copper is delivered to reproductive organs in a wheat model, brachypodium. We found that severe copper deficiency (0 or 10 nM copper) most significantly affected the development of flowers resulting in poor grain set (**Fig. 1** and **Supplemental Figure 1**). Notably, while flowers were formed in both wheat and brachypodium under low copper conditions (50 nM CuSO_4_), grain yield was severely affected (**Fig. 1E** to **H**). This “silent” effect of low copper availability on grain set could occur in crops cultivated in agricultural soils with limited copper availability underscoring the need for improving crops copper use efficiency for sustainable and environment-friendly crop production. We then show that the function of YSL3 transporter in brachypodium is essential for the transition to flowering, pollen fertility, grain yield, and quality via YSL3 function in the phloem-based copper re-distribution from mature to flag leaves and reproductive organs. This conclusion was based on the following findings. *YSL3* was expressed primarily in leaves under copper sufficient conditions and was highly upregulated by a copper deficiency in all tissues including roots, mature leaves, flag leaves and flowers but not in young leaves, possibly because the transcript level of *YLS3* in young leaves was already high (**Fig. 2**). YSL3 resided in the plasma membrane (**Supplemental Figure 2**) and the bulk of its expression was associated with the phloem in leaves and node 1 although it was also present in mesophyll and phloem parenchyma cells (**Fig. 3**). Phloem is a vascular tissue that is responsible for the translocation of nutrients including mineral element copper from source tissues such as mature leaves to sink tissues including developing flag leaves, flowers and seeds/grain [36]. Consistent with the role of YSL3 in the phloem-based copper transport from sources to sinks, copper accumulation was significantly higher in mature leaves and significantly lower in flag leaves, flowers, and grains of the *ysl3-3* mutant than of the wild-type (**Fig. 5B, C and 7A**). Because copper accumulation was significantly reduced in the phloem region in the node 1 of the *ysl3-3* mutant *vs.* wild-type (**Figure 6A, D)**, we concluded that YSL3 mediates copper loading into the phloem for subsequent distribution from source to sink tissues.

Consistent with our past studies of copper distribution in the reproductive organs of *A. thaliana* [6], the bulk of copper in florets of brachypodium was associated with anthers of stamens and ovaries of pistils (**Fig. 6C**). The inability of the *ysl3-3* mutant to deliver copper to these reproductive organs severely reduced pollen viability, germination (**Fig. 4D, E**) and significantly decreased floret fertility (**Table 1**). Importantly copper supplementation or the expression of *YSL3* cDNA rescued fertility defects of the *ysl3* mutant (**Table 1**). It is possible that the essential nature of BdYSL3-mediated copper delivery to anthers and pistils and the role of copper in pollen fertility stems from its role in maintaining metabolic functions of copper-requiring metalloenzymes, and/or for providing respiration-based energy supply for the energy-dependent reproduction processes *via* sustaining the function of the copper requiring mitochondrial cytochrome *c* oxidase complex [7, 37]. In this regard, *A. thaliana* COX11 homolog is involved in the insertion of copper into the cytochrome *c* oxidase (COX) complex during its assembly in mitochondria, is expressed in germinating pollen among other tissues and its loss of function impairs pollen germination [38]. It is noteworthy that adequate copper nutrition has been also linked to successful male fertility in mammals, including humans [39]. We also noted that copper accumulated in the stigma of pistils of wild-type plants but not of the *ysl3* mutant (**Fig. 6C**) and that the stigma of the *ysl3* mutant is less feathery compared to the wild-type (**Fig. 4F**). As the receptive portion of the gynoecium, stigma plays an important role in capturing pollen, supporting pollen germination and pollen tube guidance into the style and ovaries [40]. Finding that copper is localized to the stigma in *Brachypodium* and that loss of copper in the stigma of the *ysl3* mutant is associated with decreased fertility links stigma development and function to copper homeostasis. The role of copper in the gynoecium function is yet to be discovered.

A significant delay in transitioning to reproduction and altered inflorescence architecture as evident by nearly doubled lateral spikelet formation compared to wild-type plants (**Fig. 4 B, C** and **Table 1**) are intriguing aspects of the *ysl3* mutant phenotype. The transition from the vegetative to the reproductive stage and spikelet formation depend on the inflorescence meristem identity and determinacy, the developmental fate of axillary inflorescence meristem, which in turn depends on a variety of environmental and endogenous cues [41–44]. For example, shoot apical meristem activity in *A. thaliana* and organogenesis adapt rapidly to changes in nitrate availability in soils through the long-range cytokinin signaling [43]. Inflorescence branching and auxiliary inflorescence meristems fates in maize are regulated by sugar metabolism *via* the function of three *RAMOSA* genes [45–48]. Hormones including auxin and cytokinin have been also known to function in inflorescence architecture with auxin having a critical and conserved role in axillary meristem initiation in *A. thaliana* and maize [44, 49]. In addition to hormones, small non-coding RNA, microRNAs are implicated in developmental transitions and the regulation of inflorescence branching [49, 50]. Notably, the production of auxin and jasmonic acid is influenced by copper availability and copper deficiency stimulates the production of several miRNA families [6, 51, 52]. Considering the prominent role of copper in photosynthesis and the effect of copper homeostasis on hormone or miRNAs production, it is tempting to speculate that the defect in the internal copper distribution and delivery to flag leaves and florets in the *ysl3* mutant alters sugar metabolism, and/or miRNA and/or auxin or other hormones production resulting in delayed transition to flowering and altered inflorescence architecture. Because the timing of terminal spikelet differentiation determines the production of lateral spikelets [53, 54], it is also possible that the delayed transition to flowering observed in the *ysl3* mutant (**Fig. 4A, B**) results in increased lateral spikelets production. Although the mutation of the YSL3 orthologue in rice, OsYSL16, decreases fertility, it does not alter inflorescence architecture [23]. The distinct role of orthologous transporters may be related to distinct inflorescence architecture in rice and *Brachypodium*. The rice inflorescence, a panicle, is highly branched and is produced from multiple types of axillary meristems [44, 55]. The spikelet meristem gives rise to a single floral meristem and a single floret. By contrast, inflorescence in brachypodium is similar to its close relative wheat and is an unbranched spike where axillary meristems produced by the inflorescence meristem, develop directly into spikelets [53, 54]. Future studies will determine the specific role of YSL3 and copper in determining the inflorescence architecture of *Brachypodium*.

In addition to decreased fertility, the *ysl3-3* mutant accumulates less copper in grains and its grains are shorter, thinner and lighter than grains of wild-type, or the *ysl3* mutant expressing *YSL3* cDNA, or the mutant grown with copper supplementation (**Fig. 7**). Both grain size and weight are regulated by a complex network that integrates multiple developmental and environmental signals throughout the reproductive stage, and these processes are affected by sink and source characteristics including the size and photosynthetic capacity of source tissues and the mobilization of assimilates to the grain [56–58]. We note that the *ysl3* mutant has significantly shorter flag leaves (**Fig. 4C** and **Table 1**). Flag leaves are the most efficient functional leaves at the grain filling stage and their size and shape are among the essential traits for ideal plant-type in crop breeding programs [59, 60]. We, therefore, speculate that the decreased grain length, width, and weight in the *ysl3* mutant compared to other plant lines are caused, in part, by reduced source strength of flag leaves which, in turn, is caused by a defect in the YSL3-mediated copper distribution to flag leaves, and thus their reduced growth.

In conclusion, this study expands our understanding of the molecular mechanisms of copper transport in crop species, discovers a new avenue of copper function in establishing important agronomic traits and provides an important step towards the designing of biotechnological strategies aiming for sustainable and environmentally friendly grain yield improvement without the need for chemical fertilization in regions where poor soil quality is a major factor that limits crop productivity.

## Materials and methods

### Plant Materials and Growth Conditions

Wheat, *Triticum aestivum (cv. Bobwhite)* was used for analysis of the effect of copper on growth and reproduction. *B. distachyon* inbred line Bd21-3 regarded as wild-type [61] was used for the generation of *YSL3* mutant alleles, and transgenic plants expressing *YSL3_pro_-GUS* construct. The *ysl3*-*3* mutant allele described below was used for transformation to obtain *YSL3* complementary lines *YSL3-1* and *YSL3-2*. The generation of *YSL3* mutants and other transgenics plants is detailed in sections below. Depending on the experiment, plants were grown either in soil or hydroponically using procedures described in [27]. Briefly, after removing lamella and palea, seeds of different plant lines were surface sterilized for 10 min in a solution containing 10% bleach and 0.1% Tween 20 and then rinsed five times with deionized H_2_O. After the stratification for 24h at 4°C, seeds were sown in the water-rinsed perlite that was irrigated with ½ strength of the hydroponic solution (with or without copper). Seeds were germinated for 3 days under darkness at 24°C, then transferred to light and grown for 5 more days. The uniform seedlings were selected and transferred to soil or hydroponic solution. Hydroponic medium for both wheat and brachypodium contained 1 mM KNO_3_, 0.5 mM MgSO_4_, 1 mM KH_2_PO_4_, 1 mM Ca (NO_3_)_2_, 2.5 μM NaCl, 25 μM Fe (III)-HEDTA, 3.5 μM MnCl_2_, 0.25 μM ZnSO_4_, 0.25 μM CuSO_4_, 17.5 μM H_3_BO_3_, 0.05 μM Na_2_MoO_4_ and 0.0025 μM CoCl_2_ and this medium was replaced weekly. For achieving copper deficiency condition, plants were grown hydroponically for 3 weeks in a medium lacking copper.

Soil-grown plants were fertilized with the standard N-P-K fertilizer biweekly. To ensure that *YSL3* mutant alleles develop and produce seeds, when indicated, 25 µM CuSO_4_ was also added to the N-P-K fertilizer. In all cases, plants were grown at 24°C, 20-h-light/18°C, 4-h-dark photoperiod and a photosynthetic flux density of 150 μmol photons m^-2^ s^-1^ light produced with cool-white fluorescent bulbs supplemented by incandescent lighting and 75% relative humidity.

### Plasmid Construction

A set of plasmids was prepared for functional complementation studies, analysis of the tissue-specificity of the expression and subcellular localization of YSL3 in brachypodium, and for the generation of *YSL3* knockout plants.

The open reading frame (ORF, 2,115 bp) of *BdYSL3* without a stop codon was PCR-amplified from *Brachypodium* cDNA that was prepared from roots of plants grown hydroponically under control conditions. Three *YSL3* isoforms, Bradi5g17230.1 Bradi5g17230.2 and Bradi5g17230.3 are annotated in the *Brachypodium* genome v3.1 [62]. Because the Bradi5g17230.2 was listed as a prevailing *BdYSL3* isoform, its 2,115 bp ORF was amplified using primer pairs, *BdYSL3*-F and *BdYSL3*-R (Supplemental Table S1). The primer pairs also included att*B* sites for cloning of the PCR product by recombination into the entry *pDONR/Zeo* vector [63]. The fidelity of *BdYSL3* transcript was confirmed by sequencing. *pDONR/Zeo-BdYSL3* was then used for recombination cloning into the binary *pSAT6-N1-EGFP-Gate* [64] to fuse *BdYSL3* at the C- terminal with EGFP and place it under the control of the cauliflower mosaic virus 35S promoter. The resultant *pSAT6-N1-EGFP-Gate* with or without *BdYSL3* insert was used for the analysis of the subcellular localization of BdYSL3-EGFP in protoplasts. To study the tissue and cell-type specificity of *YSL3* expression in *Brachypodium*, a putative promoter region of *BdYSL3* (−2207 to −1 bp from the translation initiation codon) was PCR-amplified from *Brachypodium* genomic DNA using primer pairs, *BdYSL3_pro_*-F and *BdYSL3_pro_*-R (Supplemental Table S1). The amplified fragments were introduced into the p*DONR/Zeo* entry vector. After confirming the fidelity of *BdYSL3_pro_* in the p*DONR/Zeo* vector by sequencing, *BdYSL3_pro_* was transferred by recombination into the Gateway vector, *pMDC164* [65] to fuse *BdYSL3_pro_* with the bacterial *uidA* gene encoding β-glucuronidase (GUS). *pMDC164* also carries *E. coli hptII* gene conferring resistance to hygromycin for the subsequent *in planta* selection.

### The Design of CRISPR/Cas9 Constructs

To generate *YSL3* mutant alleles, we used RNA-guided DNA endonuclease system known as CRISPR/Cas9 (**<u>c</u>**lustered **<u>r</u>**egularly **i**nterspaced **<u>s</u>**hort **<u>p</u>**alindromic **<u>r</u>**epeats [CRISPR]/CRISPR-associated9 [Cas9] endonuclease) [66]. We used monocot-optimized CRISPR/Cas9 vectors that have a modular design allowing multiplexing and targeting different loci within the same gene with different single-guide (sg)RNAs simultaneously to produce larger deletions [67, 68]. Specifically, we used the module A vector, *pMOD_A1110,* that carries the wheat codon-optimized *Cas9* endonuclease gene under the control of *Zea maize Ubi* promoter, modules B and C entry vectors, *pMOD_B2518* and *pMOD_C2518*, respectively for cloning individual sgRNAs under the control of *TaU6* promoter, and the final destination vector, *pTRANS_250d* [68]. We designed CRISPR/Cas9 constructs containing two sgRNAs per construct with the intent to create larger deletions within *BdYS3* coding sequence. Thus, we designed three sgRNAs (sgRNA1, sgRNA2, sgRNA3) within the 5′ untranslated region (UTR) and the first exon of *BdYSL3,* respectively (**Supplemental Figures 3 and 4**). The targeted regions contained the CAS9-recognizing 5′-NGG protospacer adjacent motif (PAM), adjacent to the 20-bp target DNA. The lack of the off-target mutations was confirmed using the CRISPR-P 1.0 web tool (http://crispr.hzau.edu.cn/CRISPR/, [69]). The sgRNA1 and sgRNA2 were separated by 116 bp while sgRNA2 and sgRNA3 were separated by 149 bp (**Supplemental Figures 3 and 4**). sgRNA oligos were hybridized and annealed prior to cloning into the *Esp*3I site of the *pMOD_B2518* (for sgRNA1 or sgRNA2) and *pMOD_C2518* (for sgRNA2 or sgRNA3). The *pMOD_A1110* carrying *TaCas9* and two entry vectors *pMOD_B2518* and *pMOD_C2518* carrying sgRNA1 and 2, respectively or *pMOD_B2518* and *pMOD_C2518* carrying sgRNA2 and 3, respectively were combined by Golden Gate cloning [70] with the destination vector, *pTRANS_250d* to generate two *CRISPR/Cas9* destination vectors containing with sgRNA1 and sgRNA2 (sgRNAs1+2) or sgRNA2 and sgRNA3 (sgRNAs2+3); these two vectors were designated *pHS_YSL3(1+2)* and *pHS_YSL3(2+3)*, respectively.

### *Agrobacterium tumesfaciens*-mediated Transformation of *B. distachyon*

The *pMDC164* vector containing *BdYLS3_pro_-GUS*, or pSATN-EGFP-Gate vector with *BdYSL3* insert, or CRISPR/Cas 9 vectors, *pHS_YSL3(1+2)* and *pHS_YSL3(2+3)* were transformed by electroporation into *Agrobacterium tumefaciens AGL1* strain. All vectors contained the *E.coli hptII* gene conferring resistance to hygromycin for the subsequent *in planta* selection. Brachypodium transformation was done as described in [61]. Briefly, embryos were dissected from immature seeds of brachypodium and placed on callus induction medium (CIM) for 7 weeks. The formed callus was then inoculated with *A. tumefaciens* containing a construct of interest. After 3 days of co-cultivation, callus was transferred to a transformants selection medium containing 20 μg/ml hygromycin. After 6 weeks of selection, hygromycin-resistant callus was transferred to the regeneration medium. When plantlets were approximately 5 cm tall, they were transferred to clear tubes with Murasige/Skoog (MS) medium for rooting and the well-rooted plants were transplanted to soil for subsequent genotyping and seed harvesting.

### PCR Genotyping of CRISPR/Cas9 lines and Sequencing

Genomic DNA was extracted from leaves (0.1 g) using a standard cetyl-trimethyl-ammonium bromide method[71]. Twenty-five transgenic T0 lines (13 for *pHZ_YSL3 (1+2)* and 12 lines for *pHZ_YSL3(2+3)* were PCR-genotyped for the presence of deletions in the *YSL3* gene using primer pairs upstream the sgRNA1 (Genotyping-F) and downstream the sgRNA3 (Genotyping-R) (Supplemental Table S1). Deletion lines were selected by the band size and T1 generation of homozygous deletion lines was re-genotyped for the absence of *Cas9* gene using primer pairs indicated in Supplemental Table S1. Two Cas9-free deletion lines per each construct were re-genotyped for the presence of deletion using primer pairs Genotyping F and Genotyping R (**Supplemental Table S1**). PCR products were loaded onto 1% (w/v) agarose gel, excised from gel, purifying and cloned into the *pGEM-T Easy* vector (Promega) for sequencing using SP6 and T7 primers. DNA sequencing results were analyzed against the *Brachypodium* genome v3.1 [62]. Sequence alignments were done using DNAMAN software.

### Tissue- and Cell-type Specificity of *YSL3* Expression

Brachypodium Bd21-3 inbred line was transformed with *pMDC164* vector containing *BdYLS3_pro_-GUS*. Five out of the 13 independent transgenic lines (T1 generation) were used for GUS staining. Samples, collected from plants grown hydroponically with or without copper were fixed in 90% acetone on ice for 15 min. After washing thoroughly with ddH_2_O, samples were incubated at 37°C overnight in GUS staining solution containing 1 mM K_3_[Fe(CN)_6_], 1 mM 5-bromo-4-chloro-3-indolyl-β-D-glucuronide (X-Gluc), 100 mM sodium phosphate buffer (pH 7.0), 10 mM Na_2_EDTA and 0.1 % (v/v) TritonX-100[72]. After staining, samples were soaked 5 times (3-4 h each time) in 90% ethanol to remove chlorophyll that interferes with observation of the blue GUS stain. Hand-cut sections were prepared from stems using a feather double-edge razor blade. Staining patterns were analyzed using the Zeiss 2000 stereomicroscope. Images were collected using a Canon PowerShot S3 IS digital camera and a CS3IS camera adapter. Images were processed using the Adobe Photoshop software package, version 12.0.

### Functional Complementation Assays in the *Brachypodium ysl3-3* Mutant

The pSATN-EGFP-Gate vector containing the *BdYSL3* cDNA insert was transformed into the *ysl3-3* mutant allele using the described above *Agrobacterium*-mediated transformation. Two independent transgenic lines, *YSL3-1* and *YSL3-2* were selected for functional complementation assays. For plants growing hydroponically, four-week- old plants were imaged prior to tissue harvesting and biomass analysis. For plants grown in soil, days from germination to flowering were recorded for each genotype. Spike phenotypes were photographed. The floret number was estimated when seeds were ready for harvesting. The fertility was calculated as number of filled seeds per number of florets. Seed weight were measured as 1000 seeds’ weight.

### Subcellular Localization of BdYSL3

To study the subcellular localization of BdYSL3, pSATN-EGFP-Gate vector with or without *BdYSL3* cDNA insert was transfected into *A. thaliana* protoplasts by a polyethylene glycol– mediated method as described [73]. EGFP-mediated fluorescence and chlorophyll auto-fluorescence were visualized using FITC (for EGFP) or rhodamine (for chlorophyll) filter sets of the Axio Imager M2 microscope equipped with the motorized Z-drive (Zeiss). Images were obtained using the high-resolution 25 AxioCam MR Camera and processed using the Adobe Photoshop software package, version 12.0.

### RNA Extraction and RT-qPCR Analysis

Brachypodium tissues were collected from plants grown either in soil or hydroponically with or without Cu as described above. Because the expression of copper-responsive genes can be affected by the circadian rhythms [74], samples were collected at fixed time between 7 and 8 Zeitgeber hour, where the Zeitgeber hour 1 is defined as the first hour of light after the dark period. Two micrograms of total RNA extracted with the TRIzol reagent (Invitrogen) was used as a template for cDNA synthesis with the Affinity Script QPCR cDNA Synthesis Kit (Agilent Technologies). RT-qPCR and data analysis were performed as described in [75]. The expression of *ACTIN2* gene was used for data normalization. Relative expression (ΔΔCt) and fold difference (2^-ΔΔCt^) were calculated using the CFX Manager Software, version 1.5 (Bio-Rad). The gene-specific primers are listed in Supplemental Table S1.

### Elemental Analysis

Elemental analysis was performed using inductively coupled plasma mass spectrometry (ICP-MS) as described in [75, 76]. Briefly, for analysis of metal concentration in roots and young leaves, plants were grown hydroponically as described above. Root tissues were collected and desorbed in 10 mM EDTA for 5 min followed by washing in a solution of 0.3 mM BPS and 5.7 mM sodium dithionite for 10 min before rinsing three times with deionized water. For the analysis of metal concentration in flag leaves, flowers and seeds, plant lines were grown in soil. The metal concentration was determined by ICP-MS (Agilent 7700) after diluted to 10 ml with deionized water.

### Synchrotron X-ray Fluorescence (SXRF) Microscopy

Two-dimensional synchrotron x-ray fluorescence microscopy imaging the spatial distribution of copper in fresh tissues including leaves and flowers was done at the F3 station at the Cornell High Energy Synchrotron Source (CHESS). Imaging of copper distribution in nodes was done using two dimensional confocal SXRF (2D-CXRF) at beamline 5-ID (SRX) of National Synchrotron Light Source (NSLS). A detailed description of procedures is provided in the Supplementary Information.

### Pollen Viability Assays

Plants were grown in soil as described above. Pollen viability was analyzed using double staining with fluorescein diacetate and propidium iodide as described [77]. Briefly, fluorescein diacetate (2 mg/mL) was prepared in acetone and added drop by drop into 17% sucrose. Propidium iodide (1 mg/mL made in water) was diluted to a final concertation of 100 *μ*L/mL with 17% sucrose (w/v). Anthers were dissected from flowers under the stereo microscope and pollen was released by tapping into the Eppendorf tube containing fluorescein diacetate and propidium iodide solutions mixed in 1:1 ratio prior to fluorescence microscopy. Pollens were imaged under the Axio Imager M2 microscope (Zeiss, Inc) using FITC and Texas red filter sets to visualize fluorescein- and propidium iodide-mediated fluorescence. Viable pollen was stained green because live cells uptake fluorescein diacetate and convert it to fluorescein, which emits blue-green light under UV irradiation [78]. Unviable pollen was red because while propidium iodide is excluded from living cells, it labels dead cells with red-orange fluorescence under UV irradiation [79]. The number of viable and aborted pollens was counted in 10 sample microscope fields in each of three independent experiments. Images were collected with the high-resolution AxioCam MR camera and processed using the Adobe Photoshop software package, version 12.0.

### Pollen Germination Assays

Florets, collected from soil-grown plants were incubated for 1h at 4°C prior to anther dissection and pollen collection. Pollen was germinated in the medium containing 1 mM CaCl_2_, 1 mM KCl, 0.8 mM MgSO_4_, 1.6 mM H_3_BO_3_, 30 μM CaSO_4_, 0.03% casein, 0.3% MES, 10% sucrose and 12.5% polyethylene glycol [23]. Germination was scored after 3-4 h of incubation at 24 °C.

### Statistical Analysis

All the presented data are the mean values of three independent experiments. SPSS 20.0 (SPSS, Chicago, IL, USA) and JMP Pro 14.0.0 (SAS) was utilized for statistical analyses. Individual differences among means were determined using Student’s *t*-test of one-way ANOVA at a significance level of *p* < 0.05.

## Acknowledgements

We would like to thank Professor Mark Sorrells and Ellie Taagen (Cornell University) for providing *T. aestivum* seeds and for assisting in using the WinSEEDLE^TM^ of STD4800 Scanner. We would like to thank Haoyu Lin for help in phenotyping wheat; 2D-CXRF experiments used resources of the National Synchrotron Light Source II, a U.S. Department of Energy (DOE) Office of Science User Facility operated for the DOE Office of Science by Brookhaven National Laboratory under Contract No. DE-SC0012704. H.S. was supported by National Natural Science Foundation of China (# 31301349, 30870154, 30901052, 30900087) award to Y.Z. and The Schwartz Research Fund for Women in Life Sciences awarded to O.K.V, This study was funded by USDA/NIFA NYC-125542, NSF-IOS #1656321, The Schwartz Research Fund for Women in Life Sciences awarded to O.K.V., and CRDF-GLOBAL U.S.-Ukraine Competition OISE-9531011, awarded to O.K.V., O.I.T and N.D.R.

## Author Contributions

O.K.V. designed experiments with H.S.; H.S., Y.J., M.R.I, J.-C.C., P.M., T.Z., and Y.K., performed experiments; T.D. assisted in ICP-MS analysis; R.H., L.S. and A.W. facilitated the SXRF experiments at CHESS and NSLSII (Brookhaven National labs). Y.Z., O.I.T., N.D.R. contributed to discussion on the manuscript. The manuscript was written by O.K.V and H.S. All authors contributed constructive comments on the manuscript.

## Figure legends

**Supplemental Figure 1. Copper deficiency affects flower development and reduces tiller and head number in wheat and brachypodium.** Plants were grown hydroponically under indicated copper concentrations. (**A)** and (**B**) show a representative image of spike appearance under different copper concentrations in wheat or brachypodium, respectively. (**C)** to (**F)** show the number of tillers and heads formed per plant in wheat (**C, D**) or brachypodium (**E, F**). Statistical analysis in (**C)** to (**F)** was done with one-way ANOVA in the JMP Pro 14 software package; comparison of the means for each pair was done using Student’s *t*-test. **C** to **F** show mean values ± S.E. from the analysis of 4 (wheat) to 6 (brachypodium) independently grown plants from one out of two (wheat) and three (brachypodium) independent experiments. Levels not connected by same letter are significantly different (*p* < 0.05).

**Supplemental Figure 2. BdYSL3 localizes to the plasma membrane in *A. thaliana* protoplast**. *BdYSL3* fused with EGFP at the C-termini (**A**) or EGFP-expressing empty vector (**B**) were transfected into protoplasts prepared from *A. thaliana* mesophyll cells. Shown are bright field image (**Bf**) of transfected protoplasts, YSL3-EGFP (**YSL3**)-mediated fluorescence, EGFP (**EGFP**) - mediated fluorescence and chlorophyll autofluorescence (**Chl**) fluorescence. Overlay images were created to show that YSL3-EGFP-mediated fluorescence does not co-localize with the chlorophyll-mediated fluorescence. Scale bar = 5 µm.

**Supplemental Figure 3. Generation of CRISPR/Cas9 mutant alleles for *YSL3*.** (**A**) The schematic illustration of *BdYSL3* (Bradi5g17230.2) gene structure. Black boxes show exons, gray line show introns and 5′and 3′UTRs. The close-up marked by a dashed box shows *YSL3* region targeted by sgRNAs (gRNA1, gRNA2, gRNA3), shown as red bars. The expected deletion sizes are indicated. Blue arrows show the relative position of the forward (F) and revers (R) genotyping primers. (**B**) PCR-based genotyping of four T1-generation of CRISPR/Cas9 transgenic lines of which lines 1 and 2 are from the transformation with a construct containing sgRNA2 and 3 (gRNA2+3) while lines 3 and 4 are from the transformation with a construct containing sgRNA1 and 2 (gRNA1+2). (**C**) shows results of PCR-based genotyping using primers against *Cas9* in the *CRISPR/Cas9* construct or against brachypodium *Actin*. Genomic DNA was isolated from a wild-type plant (**Wt**), T1 plants for lines 1 to 4 (1, 2, 3, 4) or a T0 plant (T0). The latter was used as a control for *Cas9*. (**D**) *ysl3* deletion lines and untransformed wild-type plants were grown hydroponically for 3 weeks with 0.25 µM CuSO_4_. (**E**) Sequencing results of plant transformed with sgRNA 1 and 2 (lines 3 and 4) show 122 and 123 bp deletion in plants. These mutant lines are designated *ysl3-1* and *ysl3-2*, respectively. Sequencing results of plant transformed with sgRNA 2 and 3 (lines 1 and 2) show 182 bp deletion. Both lines represent the same allele which is designated as *ysl3-3*.

**Supplemental Figure 4.** The genomic DNA sequence of *YSL3* that was used for designing sgRNAs. The position of sgRNAs and sequencing primers are indicated in bold and blue font, respectively. The Protospacer Adjacent Motif (PAM) sequence is shown in red.

**Supplemental Figure 5. RT-qPCR analysis of *YSL3* transcript abundance in roots and leaves of *Brachypodium* wild-type (*Wt*), the *ysl3-3* mutant (*ysl3*) and two *ysl3-3* mutant transgenic lines expressing the *BdYSL3* cDNA (*YSL3-1; YSL3-2).*** Plants were grown hydroponically in the presence of 0.25 µM CuSO_4_. Roots and shoots were collected from the 4-week-old plants. *YSL3* expression in different plant lines are shown relative to its expression in leaves of Wt that was designated as 1. Error bars indicate S.E. (n = 3 pools of plant tissues each collected from 3 plants. Levels not connected by same letter are significantly different (*p* < 0.05).

**Supplemental Figure 6. Y*S*L3 is essential for the normal growth of *Brachypodium* under copper deficiency**. (**A)**. Wild-type plants (*Wt*), the *ysl3-3* mutant (*ysl3*) and two *ysl3-3* mutant transgenic lines expressing the *BdYSL3* cDNA (*YSL3-1* and *YSL3-2*) were grown hydroponically with the indicated concentrations of CuSO_4_. Representative photos of nine plants analyzed per each line were captured after three weeks of growth. (**B)**. Representative images of top (youngest) leaves collected from plants grown hydroponically for 3 weeks with the indicated concentrations of CuSO_4_. A white arrow points to a leaf with curled margins. (**C**) shows the height, (**D, E**) shows the biomass of the *Brachypodium* wild-type plants (*Wt*) *ysl3-3* mutant (*ysl3*), and two *ysl3-3* mutant transgenic lines expressing the *BdYSL3* cDNA (*YSL3-1* and *YSL3-2*) grown hydroponically for 3 weeks with the indicated concentrations of CuSO_4_. **C** to **E** show mean values of 6 independently grown plants from three independent experiments, error bars show S.E. Levels not connected by same letter are significantly different (*p* < 0.05).

**Supplemental Figure 7. The *ysl3-3* mutant is not sensitive to iron, manganese or zinc deficiency**. Indicated plant lines were germinated in perlite for 1 week, then a subset of plants was transferred to either to a complete hydroponic medium containing 1 µM CuSO_4_ to allow growth of the *ysl3* mutant (Control), or the same medium that lack either manganese (-Mn) or iron (-Fe), or zinc (-Zn). Plants were photographed after 3 weeks. A representative image from three independent experiments is shown.

